# Pluronic gel-based burrowing assay for rapid assessment of neuromuscular health in *C. elegans*

**DOI:** 10.1101/632083

**Authors:** Leila Lesanpezeshki, Jennifer E. Hewitt, Ricardo Laranjeiro, Adam Antebi, Monica Driscoll, Nathaniel J. Szewczyk, Jerzy Blawzdziewicz, Carla M.R. Lacerda, Siva A. Vanapalli

**Affiliations:** Department of Chemical Engineering, Texas Tech University, Lubbock, TX, USA; Department of Molecular Biology and Biochemistry, Rutgers, The State University of New Jersey, Piscataway, NJ, USA; Department of Molecular Genetics of Ageing, Max Planck Institute for Biology of Ageing, and Cologne Excellence Cluster on Cellular Stress Responses in Aging-Associated Diseases (CECAD), University of Cologne, Cologne, Germany; MRC/Arthritis Research UK Centre for Musculoskeletal Ageing Research, University of Nottingham, United Kingdom & National Institute for Health Research Nottingham Biomedical Research Centre, Derby, UK; Department of Mechanical Engineering, Texas Tech University, Lubbock, TX, USA

## Abstract

Whole-organism phenotypic assays are central to the assessment of neuromuscular function and health in model organisms such as the nematode *C. elegans*. In this study, we report a new assay format for engaging *C. elegans* in burrowing that enables rapid assessment of nematode neuromuscular health. In contrast to agar environments that pose specific drawbacks for characterization of *C. elegans* burrowing ability, here we use the optically transparent and biocompatible Pluronic F-127 gel that transitions from liquid to gel at room temperature, enabling convenient and safe handling of animals. The burrowing assay methodology involves loading animals at the bottom of well plates, casting a liquid-phase of Pluronic on top that solidifies via a modest temperature upshift, enticing animals to reach the surface via chemotaxis to food, and quantifying the relative success animals have in reaching the chemoattractant. We study the influence of Pluronic concentration, gel height and chemoattractant choice to optimize assay performance. To demonstrate the simplicity of the assay workflow and versatility, we show its novel application in multiple areas including (i) evaluating muscle mutants with defects in dense bodies and/or M-lines (*pfn-3, atn-1, uig-1, dyc-1, zyx-1, unc-95* and *tln-1*), (ii) tuning assay conditions to reveal changes in the mutant *gei- 8*, (iii) sorting of fast burrowers in a genetically-uniform wild-type population for later quantitation of their distinct muscle gene expression, and (iv) testing proteotoxic animal models of Huntington and Parkinson’s disease. Results from our studies show that stimulating animals to navigate in a dense environment that offers mechanical resistance to three-dimensional locomotion challenges the neuromuscular system in a manner distinct from standard crawling and thrashing assays. Our simple and high throughput burrowing assay can provide insight into molecular mechanisms for maintenance of neuromuscular health and facilitate screening for therapeutic targets.

## I. Introduction

The millimeter-long round worm *Caenorhabditis elegans* is an excellent model for investigating conserved molecular mechanisms regulating neuromuscular health and its decline with age [1]. Additional benefits of this model organism include 60% homology with human genes, a short life cycle, 95 body wall muscle cells similar to vertebrate muscle, and 302 mapped neurons [1–3]. Studies in *C. elegans* can potentially lead to insights into maintenance of neuromuscular health [4], impacting a wide range of human disorders ranging from muscular dystrophies [5] to neurodegenerative diseases [6].

Whole-organism assays are an essential aspect of scoring neuromuscular health and studying mechanisms regulating neuromuscular function. Assays typically involve investigating *C. elegans* locomotion on agar plates [7], where animals crawl in two-dimensions (2D). Alternatively, thrashing assays that score animals swimming locomotion [8] or microfluidic pillar environments that measure muscle forces have been used [9, 10]. Additionally, sophisticated computer-vision analysis can be employed to further reveal detailed aspects of *C. elegans* locomotion and maneuverability in these different environments [11–13].

Although existing methods have revealed important aspects of neuromuscular control of animal behavior [14], assay culture conditions are typically not the best representation of *C. elegans* natural habitat, in which animals burrow in three-dimensions in soil, rotten fruit, and fluid drops [15]. Indeed, burrowing environments to mimic this natural habitat have been designed to better reflect true behavioral conditions [16–20].

3D locomotion in *C. elegans* is distinct from 2D locomotion, because the neuromuscular control system needs to enable body twisting to generate out-of-plane motion. These roll maneuvers are essential for the animal to make turns and navigate in 3D [21]. Evidence is emerging that 3D locomotory studies can provide information on *C. elegans* genetics and neuromuscular aspects that are not evident in 2D measures [16, 17]. For example, in mutants modeling muscular dystrophy, differences observed in their 2D crawling performance were minor compared to an obvious burrowing defect [16]. As a result, engaging animals in burrowing environments and scoring their locomotory prowess might provide the sensitive dynamic range of outcomes to reveal previously undiscernible phenotypes.

Burrowing capacity has been examined in agar-filled pipettes by stimulating animals to move towards the food source present at one end of the pipette [16, 18, 22]. A limitation of this burrowing assay is that the methodology requires time-consuming steps including drilling holes in the pipette (later used for loading animals and food source), followed by filling the pipettes uniformly with hot agar and waiting for them to cool. Additional challenges include injection of tens of animals into a localized spot (which may result in animal loss or damage), and visualization difficulty due to the translucency of agar and refraction at cylindrical surfaces.

Despite the above limitations, the basic approach and the burrowing assay are highly valuable - stimulating animals to burrow in a dense, 3D environment offers a way to challenge the neuromuscular system that is distinct from swimming or 2D crawling that may well approximate a more natural environment than either swimming or crawling. Here we report on a novel design for burrowing evaluation that addresses the limitations of the currently described agar-based burrowing assay and enables parallel evaluation of burrowing performance of wild-type and mutant *C. elegans*. The method utilizes well plates and Pluronic F-127—an optically transparent biocompatible hydrogel that undergoes a sol-gel transition in a temperature range that is safe for handling *C. elegans*. We tested the influence of system parameters including gel concentration, height, and chemoattractant type to optimize the assay performance. In addition, we evaluate animal distribution and behavior during burrowing.

To demonstrate the flexibility and power of the assay, we show its suitability for diverse applications including (i) evaluation of muscle-defective mutants, (ii) tuning assay conditions to increase phenotypic distinction of a mitochondrial mutant, (iii) separation and recovery of fast/slow burrowers for differential gene expression analysis, and (iv) testing disease models of protein aggregation for locomotory impairment. We anticipate that this new and simple assay should provide insights into molecular mechanisms regulating maintenance of neuromuscular health, facilitate screening for pharmacological interventions, and create a path to novel discovery of therapeutic targets.

## II. Results

### A. Basic methodology of Pluronic-based burrowing assay

To create a 3D environment for burrowing, a medium that confers no adverse effect on animal health is essential. It is a standard practice to culture *C. elegans* on agar (in the form of Nematode Growth Media (NGM)) plates, and thus studies so far have focused on using agar as the burrowing medium [16, 18, 20]. Although agar is an animal-friendly medium, *C. elegans* cannot tolerate the high temperatures (> 32-40 °C) of liquid agar, and therefore execution of a burrowing assay requires forcible injection of animals into the agar after it has cooled down and solidified. The mechanical stress during injection may not only rupture the gel, but also can result in animal injury and loss as a high density of animals are introduced into a localized area. Moreover, in the published burrowing method [16, 18], animals are not easy to visualize due to the translucency of agar and refraction at the cylindrical surfaces of glass pipettes that are used.

Here, we introduce Pluronic F-127 (PF-127), a biocompatible thermoreversible hydrogel, as a novel 3D gel environment for a burrowing assay. PF-127 is a copolymer consisting of a hydrophobic propylene oxide block in the center and flanked by two hydrophilic ethylene oxide blocks. PF-127 has become popular for immobilizing *C. elegans* and for high resolution imaging due to its excellent optical transparency [23–27] and its highly favorable range for solid/fluid transitions. Furthermore, a recent study [26] showed that *C. elegans* can tolerate continuous exposure to PF-127 up to 4 hours, making it a safe medium for *C. elegans* assays of short duration.

Our current burrowing assay format uses 12-well plates, which allows for assessment of four treatments each with three replicates per plate (Fig. 1a). The optimized protocol involves loading a liquid drop (20 - 30 μL) of 26 % w/w PF-127 at 14°C at the bottom of the well, which forms a thin (< 1 mm) gel layer within a couple of minutes at room temperature (20°C ± 2°C). About 30 hand-picked animals taken from culture plates are introduced on this bottom layer. We note that since the thin gel layer is equilibrated to room temperature before adding animals, they are not exposed to significant temperature shock. Next, a second viscous liquid layer of 26 % w/w PF-127 is cast on top of the loaded animals to achieve a desired thickness, which gels within 5 mins at room temperature. A chemoattractant is then placed on top of the gelled surface and the number of animals reaching the top is counted every 15 minutes for a minimum of 2 hours (See supplementary video S1).

**Figure 1.**
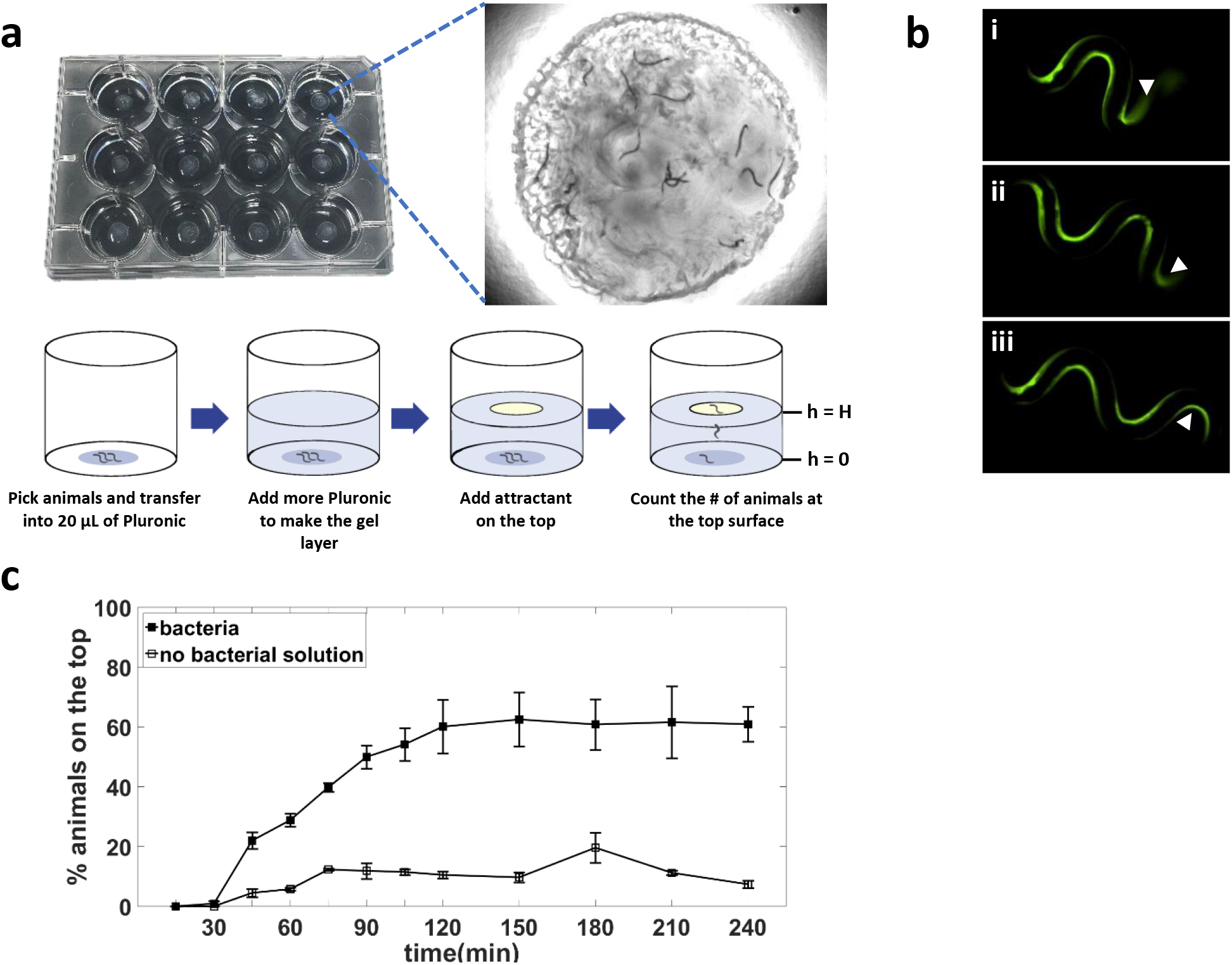
Basic principle of Pluronic (PF-127) gel-based burrowing assay. (a) Burrowing assay is conducted in a 12 well-plate; in each well animals are stimulated to move through a gel towards a food source placed at the top. The inset shows the nematodes that have successfully burrowed and reached the food source (*E. coli*) at the top of the gel (See also supplementary video S1). (b) Calcium imaging shows the muscle contractions as the nematode is burrowing in the PF-127 gel. Strain is HBR4: goeIs3 HBR4: goeIs3[*Pmyo-3*::GCaMP3.35::*unc-54*-3’utr, *unc- 119*] expressing the calcium indicator GCaMP3 in body wall muscles. Images are taken with 5s intervals. Arrow heads point to the tail that appears faded in i and ii due to the 3D locomotion. In all three images, the head is on the left. (c) Burrowing performance of day 1 adults wild-type animals in the presence and absence of *E. coli* bacteria in 26 % w/w PF-F127. Gel thickness, H = 0.9 cm. N = 39 and 35 animals in the presence and absence of bacteria, respectively.

We tested wild-type animals in the presence and absence of chemotactic stimulus, *E. coli* OP50 (the animal’s standard food source) (Fig. 1b). We observed that in the presence of the stimulus, the percentage of animals reaching the top surface increased with time, saturating at ≈ 60% after 2 hours. In contrast, without any chemotactic stimulus added to the gel surface, the percentage of animals reaching the top remained < 20%. Since the assay engages the animals to burrow from the bottom toward the top, going against gravity, we also tested whether gravitactic stimulus influences the burrowing performance. We conducted a set of experiments where the well-plate was turned upside down after introduction of the chemoattractant, and found no considerable impact on the percentage of the animals reaching the gel surface (Fig. S1). Thus, the overall direction of burrowing with respect to gravity does not influence burrowing performance, and chemotactic stimulus is essential for animals to burrow efficiently towards the surface and engage in 3D locomotion.

To confirm that the animals indeed actuate their muscles and undergo 3D out-of-plane motion by twisting their body when they navigate in the Pluronic gel environment, we used transgenic animals (P_*myo-3*_GCaMP3.35) expressing the calcium sensor GCaMP3.35 in body wall muscles[28] (Fig. 1b, also see supplementary video S2). As muscle contracts, calcium ions are released from the sarcoplasmic reticulum into the cytosol, where they interact with the sensor and generate signals associated with calcium elevation [28–30]. During burrowing, we observe that the muscle cells appear brighter on the side where the body is contracting while the other side with relaxed muscles is dark due to the momentarily low calcium activity. In Fig. 1b, the arrows point to the tail that appears faded in the first two images due to the animal’s 3D posture; however, in the last image the entire body emerges in the focal plane of the microscope. Thus, the animals undergo 3D locomotion in the PF-127 gel environment.

### B. Optimization of burrowing assay parameters

To determine optimal performance for the burrowing assay, we considered several parameters. Animal performance during burrowing is expected to depend on the mechanical resistance offered by the gel. This resistance can be controlled by the gel’s mechanical properties, which in turn is governed by PF-127 concentration. In addition, the gel height dictates how far animals need to burrow. Finally, the chemotactic response can be sensitive to the type of chemoattractant used to stimulate the animals to burrow. In this section, we therefore study the influence of gel concentration, gel height and chemoattractant choice on the burrowing performance of wild-type animals.

#### Gel concentration

Previous studies show that the sol-gel transition temperature decreases with increasing Pluronic concentration [23, 31]. For example, it has been reported that at 19% w/v the sol-gel transition occurs at 24 °C, whereas at 30% w/v, the sol-gel transition occurs at 12 °C [23]. To identify the optimal concentration range, our criteria were: (i) the sol-gel transition must occur below room temperature; (ii) the liquid should not be too viscous to handle below the sol-gel transition because Pluronic solution needs to be transferred in liquid phase to the well-plates; (iii) the handling temperature should not be far below room temperature (otherwise the gel takes too long to equilibrate to room temperature). Considering these factors, the useful PF-127 concentration range to test was between 24 to 30% w/v. For transferring the solution in liquid phase, we chose 10-14 °C, temperatures right below the sol-gel transition for this concentration range [23]. Our initial trials showed that the second layer gels in less than 5 minutes under these conditions.

Fig. 2a shows the burrowing data for PF-127 concentrations of 24, 26, 28, and 30 % w/w at a set gel height of 0.7 cm. The transfer temperature used here was 14 °C for 24, 26, and 28 % w/w, and 10 °C for 30 % w/w. At the 2-hr timepoint, we find that the percentage of animals reaching the top surface for 24, 26, 28, and 30 % w/w is 85%, 77%, 64%, and 40% respectively. Of note, we found the burrowing percentage plateaued after ≈ 60 min and 90 min for 24 and 26 % w/w respectively, but did not saturate within the 2 hrs for 28 and 30 % w/w.

**Figure 2.**
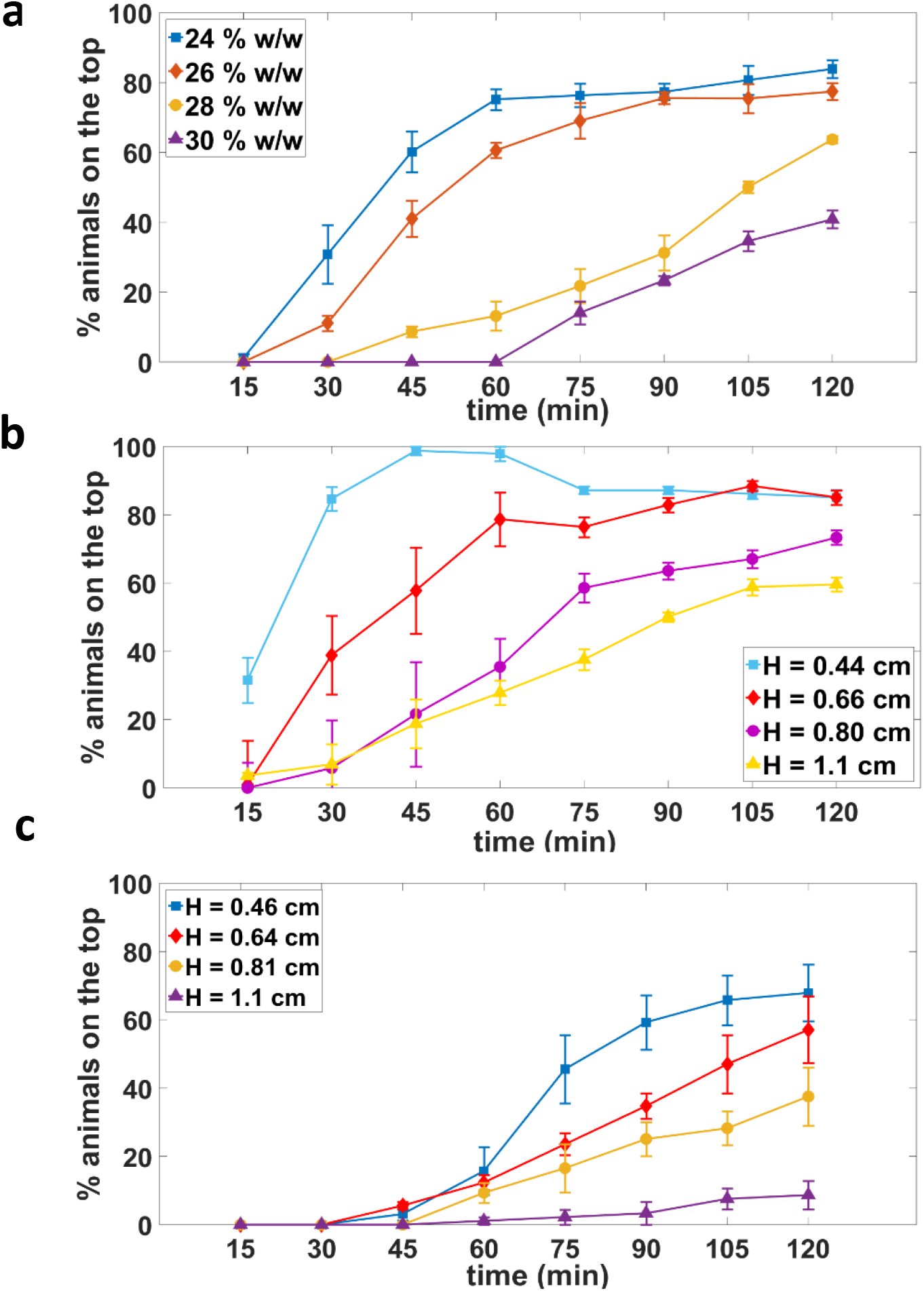
Effect of Pluronic gel concentration and height on burrowing performance of wild-type animals. Burrowing performance in (a) four different concentrations of PF-127. N = 31, 25, 31 and 34 animals for 24, 26, 28 and 30 % w/w gels respectively; H=0.7 cm. (b) 26 % w/w PF-127 at four different gel heights. N = 28, 33, 23, 28 animals for H = 0.44, 0.66, 0.80 and 1.1 cm, respectively. (c) 30 % w/w PF-127 at four different gel heights. N = 31, 30, 32 and 31 animals for H = 0.46, 0.64, 0.81 and 1.1 cm, respectively. 100 mg/mL *E. coli* was used as an attractant.

Our observation that fewer animals reach the surface as the Pluronic concentration increases, suggests that the mechanical resistance offered by the gel is an important factor determining burrowing performance. Indeed, the measured elastic modulus and yield stress of the gel increase when the PF-127 concentration increases (Fig. S2). The mechanical resistance can come from the nematode trying to overcome the yield stress to carve a hole in the gel in order to move. Also, the viscous friction along the nematode body could contribute to resistance to motion in 3D.

In sum, our results show that the mechanical resistance to burrowing can be easily tuned by modulating the PF-127 concentration. From an assay perspective, we chose PF-127 concentration of 26 % w/w as the standard condition since the challenge for animal burrowing is intermediate, and the solution gels quicker than the 24 % w/w.

#### Gel height

With respect to gel height H, we anticipated that burrowing performance should decrease with an increase in gel height. In Fig. 2b, we indeed observe this trend when we tested gel heights of H = 0.66, 0.80, and 1.1 cm at 26 % w/w PF-127 concentration. The final 2-hr burrowing percentage decreases from 85% to 60% when using gel heights of H = 0.66 and 1.1 cm, respectively. Interestingly, when we tested H = 0.44 cm, the burrowing percentage reached close to 100% within the first hour and subsequently declined as some of the animals burrowed back into the gel.

We also tested the influence of gel height at 30 % w/w of PF-127 (Fig. 2c). The same decreasing trend of burrowing performance with an increase in gel height was observed. Even at the lowest height of 0.46 cm, only 68% of the animals made it to the top by the end of 2 hours, which is considerably lower than 85% in 26 % w/w at the same height (Fig. 2b). The burrowing performance was the worst in the highest gel thickness we tested (1.1 cm), as less than 10% of the animals could reach the surface.

For the rest of study, we conducted burrowing assays with PF-127 concentration of 26 % w/w and gel height of approximately 0.7 cm unless otherwise noted. Under these conditions, we found that the burrowing percentage at the end of two hours was 70%-80% for wild-type animals.

#### Chemoattractant

The main premise of the burrowing assay is to stimulate the animals to burrow from the bottom of the well plate to the top surface of the gel. *Escherichia coli* OP50, the typical *C. elegans* food source, can act as a chemoattractant to these animals [32]. Odorant chemical compounds such as isoamyl alcohol and diacetyl are also known to be attractive to *C. elegans* (Fig. S3) [33]. To achieve a reliable burrowing performance, a chemical stimulus with a maintained level of animal attraction over the assay time should be utilized. We therefore evaluated the burrowing performance of wild-type animals stimulated with *E. coli* in salt solution or odorant chemical compounds.

We tested four concentrations of *E. coli* varying from 100 mg/mL concentrated in liquid NGM to 0 mg/mL separately. The 2-hour burrowing performance gradually diminished as more diluted attractants were utilized in each assay (Fig. 3a). Interestingly, liquid NGM without bacteria also appeared to be attractive to the animals, which could be related to salt and/or amino acid taxis [34]. Although 1% diacetyl and 1% isoamyl alcohol in ethanol are clearly attractive and elicit strong burrow outcomes in 90 minutes, animals burrow back into the gel thereafter, so attraction is not maintained (Fig. 3b). On the contrary, with *E. coli* as a stimulant, the animals tend to stay and feed on the bacteria layer when they reach the top surface of the gel, making it easier to sort the animals as they reach the surface. Taken together, our data suggest that *E. coli* at a concentration of 100 mg/mL liquid NGM is an efficacious and near optimal attractant for our burrowing assay.

**Figure 3.**
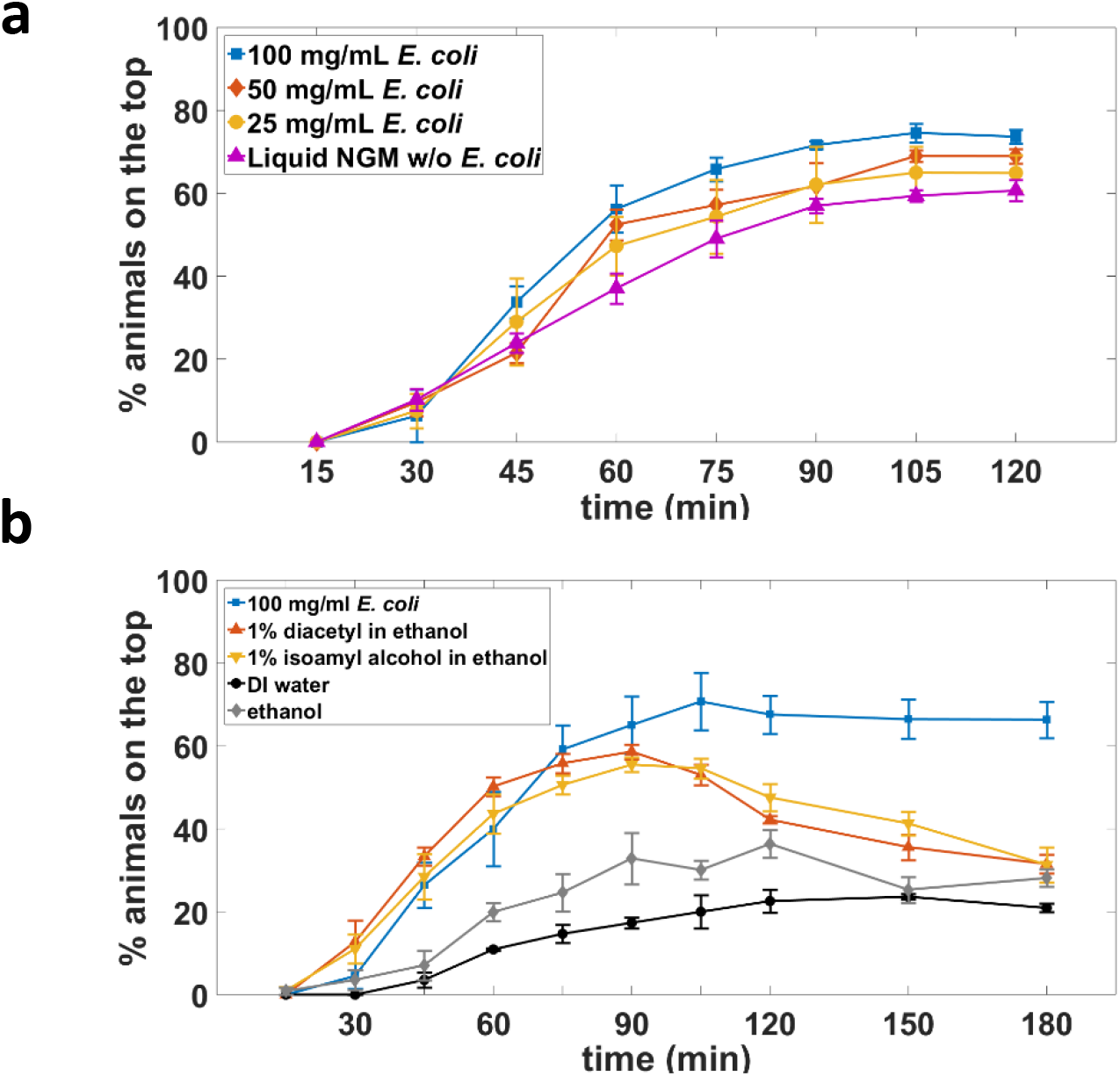
Chemoattractant choice influences the burrowing performance in wild-type animals. (a) Influence of different *E. coli* concentration on burrowing success. Assay conditions are 26 % w/w PF-127 and H =0.7 cm. N = 35, 28, 31 and 30 animals for *E. coli* concentrations of 100, 50, 25 and 0 mg/mL respectively. (b) Effect of different chemoattractants on burrowing performance. Assay conditions are 26 % w/w PF-127 and H = 0.75 cm. N = 38, 34, 33, 37 and 37 animals for 100 mg/mL *E. coli*, 1% diacetyl, 1% isoamyl alcohol, water and ethanol respectively.

### C. Animal locomotory and behavioral analysis during burrowing

To compare the locomotory characteristics in PF-127 gel with swimming and crawling, we examined *C. elegans* locomotion in a 26 % w/w Pluronic gel with a layer thickness of approximately 1 mm. This thin-gel configuration restricted the significant out-of-plane motion typically observed in our standard burrowing assay and facilitated quantitative characterization of velocity and undulatory frequency at an individual animal level. We find that the velocity ranges from 0.5 – 1.9 mm/min for individuals with a population mean of 1.14 mm/min (Fig. 4a). These velocity scores are lower than typical values reported for crawling [16, 35]. Likewise, the undulatory frequency range is 0.03 – 0.1 Hz (Fig. 4b) with a population mean of 0.07 Hz, which is also lower than the values reported for crawling and swimming [16, 35]. The diminished locomotory metrics in the Pluronic gel suggest that the burrowing environment poses significant mechanical resistance for animal locomotion. We also observed that in the thin-gel environment, animals spend 81% of their time moving forward, 15% of their time moving backward, and only 4% of their time pausing (Fig. S3). Overall, although the gel environment does restrict some locomotion metrics compared to swimming and crawling environments, animals still successfully and continually navigate through the dense gel.

**Figure 4.**
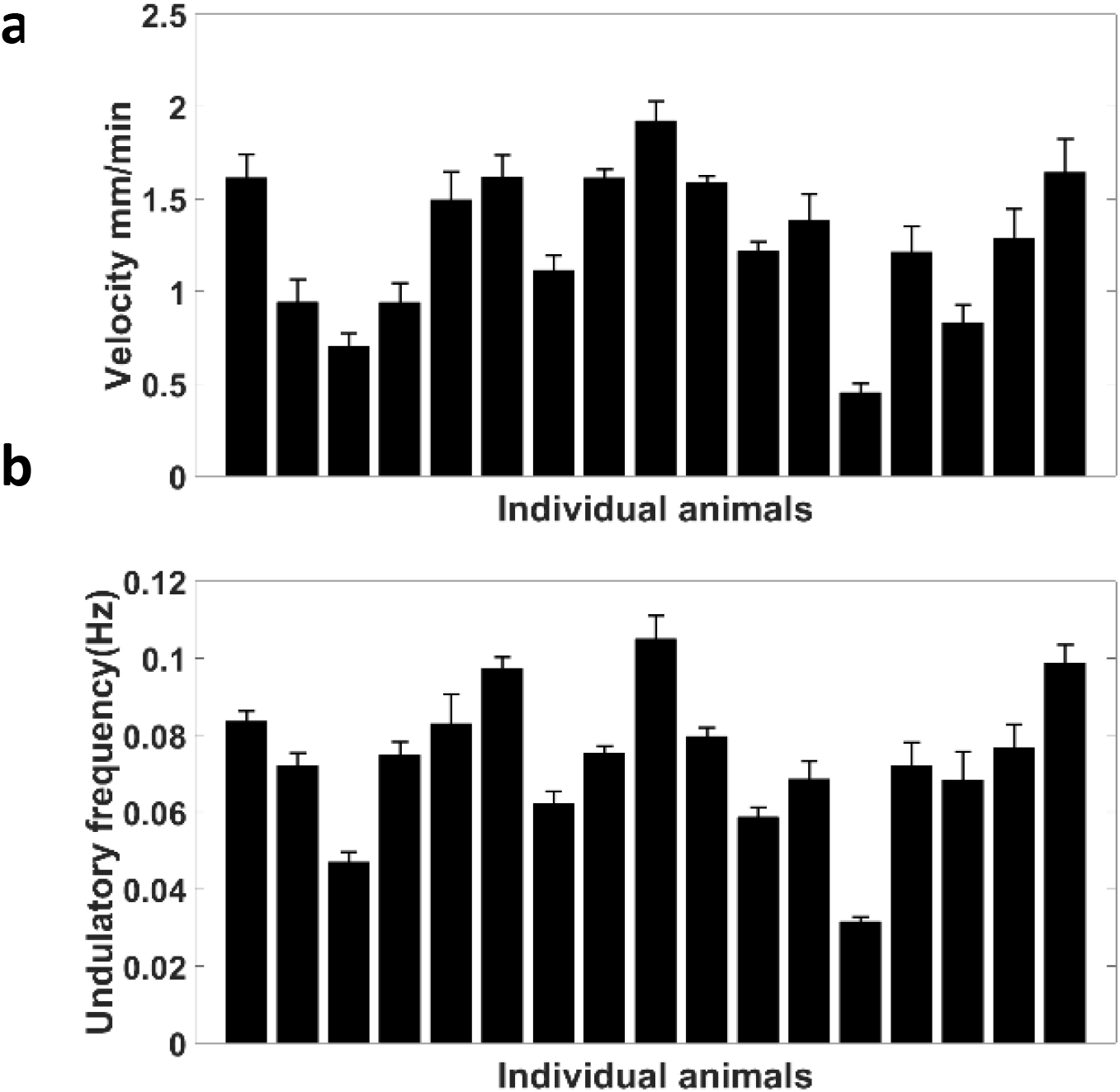
Characterization of wild-type animal locomotion in a thin Pluronic gel layer. (a) Individual animal velocity obtained by tracking the body centroid every 15 seconds. The average burrowing velocity is 1.14 mm/min. Error bar is standard error of mean calculated from 8 time intervals per individual animal. (b) Undulatory frequency of individuals obtained from the time required to complete one sinusoid body movement. The average undulatory frequency is 0.07 Hz. Error bar is standard error of mean calculated from 8 replicates per individual animal. In (a) and (b) no food source was used and the assay conditions were 26 % w/w PF-127 and H ≈ 1 mm.

To further evaluate whether the stimulated burrowing behavior contains long resting phases during their journey to reach the surface of the gel, we assessed the distribution of animals across the gel height in 15 min intervals. We monitored the fraction of animals present in consecutive layers divided into thirds, with the fourth category being the top surface. Fig. 5a demonstrates that the animals do not spend much time burrowing in the middle layers, meaning that once they sense the attractant on top, they engage in burrowing to reach the attractant. As the percentage of animals decreases in the loading area, the percentage of animals on the surface consequently increases. If there is no attractant on the top surface to stimulate the animals, animals prefer to not move much farther vertically from their initial location (Fig. 5b). Thus, we do not observe long resting phases (i.e., on the scale of several minutes) when animals are stimulated to burrow.

**Figure 5.**
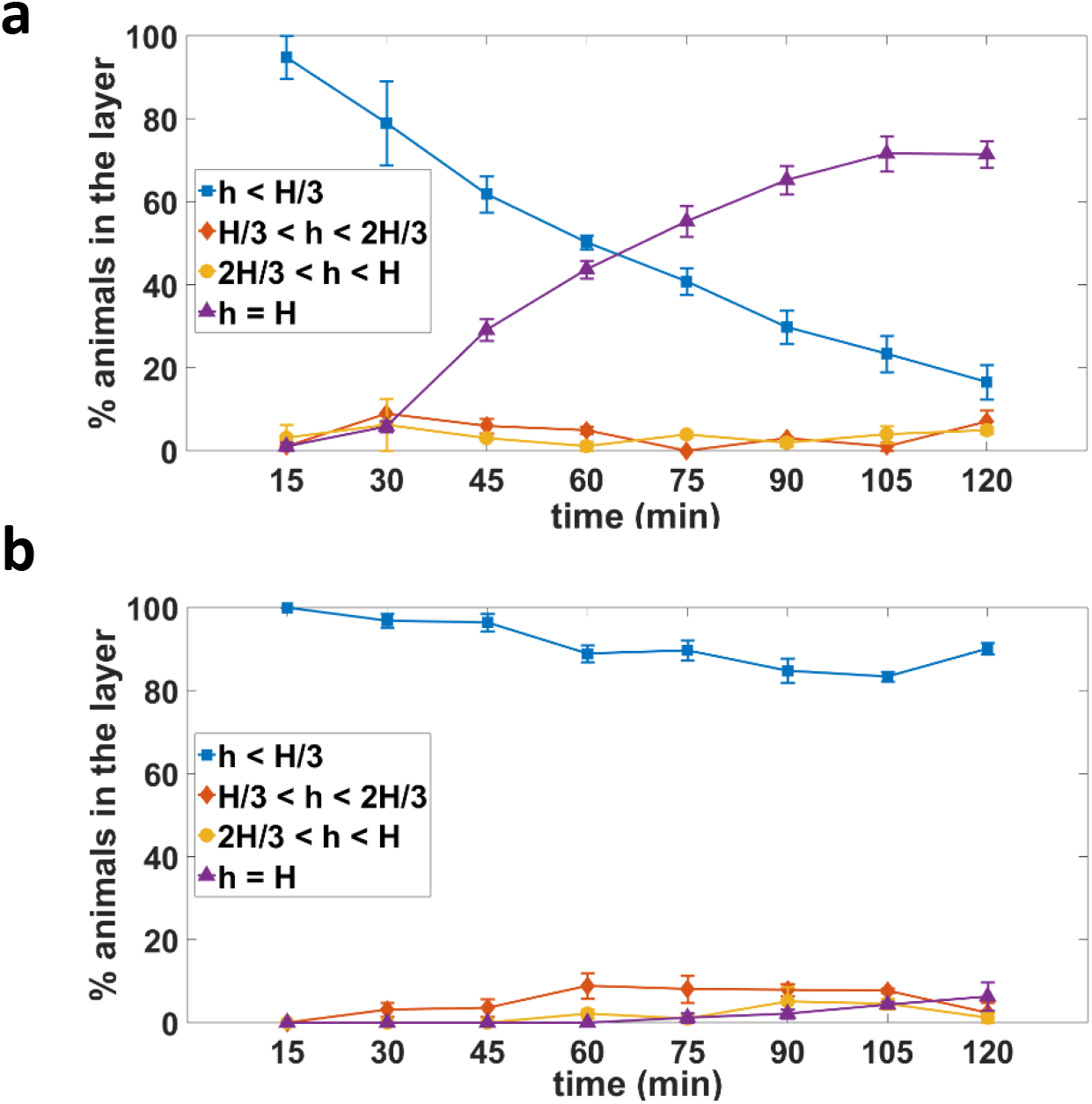
Distribution of wild-type animals during stimulated burrowing. Animal distribution in different vertical layers of the gel (a) in the presence of food source with h = H being the top where the attractant has placed (See Fig. 1a), N = 33 animals and (b) in the absence of food source, N = 30 animals. Assay conditions were 26 % w/w PF-127, H = 0.9 cm and 100 mg/mL *E. coli*.

### D. Pluronic-based burrowing assay is well suited for diverse *C. elegans* applications

Our results show that Pluronic gel as a medium for *C. elegans* burrowing offers several unique advantages including ease of manipulation of animals, easy adjustment of assay conditions to modulate burrowing performance, and the capacity to run assay conditions in parallel in multi-well plates. In this section, we harness these capabilities to demonstrate that burrowing can be used in diverse applications. None of these demonstrative applications and the results thereof have been shown by prior burrowing studies [16–20]. Since our intent is to demonstrate the power of the assay platform we designed, we consider our results in each of these applications as laying foundation for future in-depth investigations.

#### i. Mutants with muscle structural defects are deficient in burrowing

Given that the burrowing assay challenges the locomotory ability of *C. elegans* and that calcium imaging shows muscle contractions, we assessed the suitability and sensitivity of the assay to score mutants with known genetic defects in muscle. For this analysis, we chose mutations in genes encoding proteins localized to dense bodies and M-lines, the two main structural components responsible for assembly and maintenance of the sarcomeres that generate force in *C. elegans* muscle. Dense body and M-line proteins are also responsible for force transduction from contractile apparatus to hypodermis and cuticle to assist with movement [36] (Fig. 6a). Individual gene knockdown of members of these multiprotein complexes has revealed that they are both critical for muscle structure and force transmission, and required for maintaining muscle protein homeostasis and muscle mitochondrial function [37].

**Figure 6.**
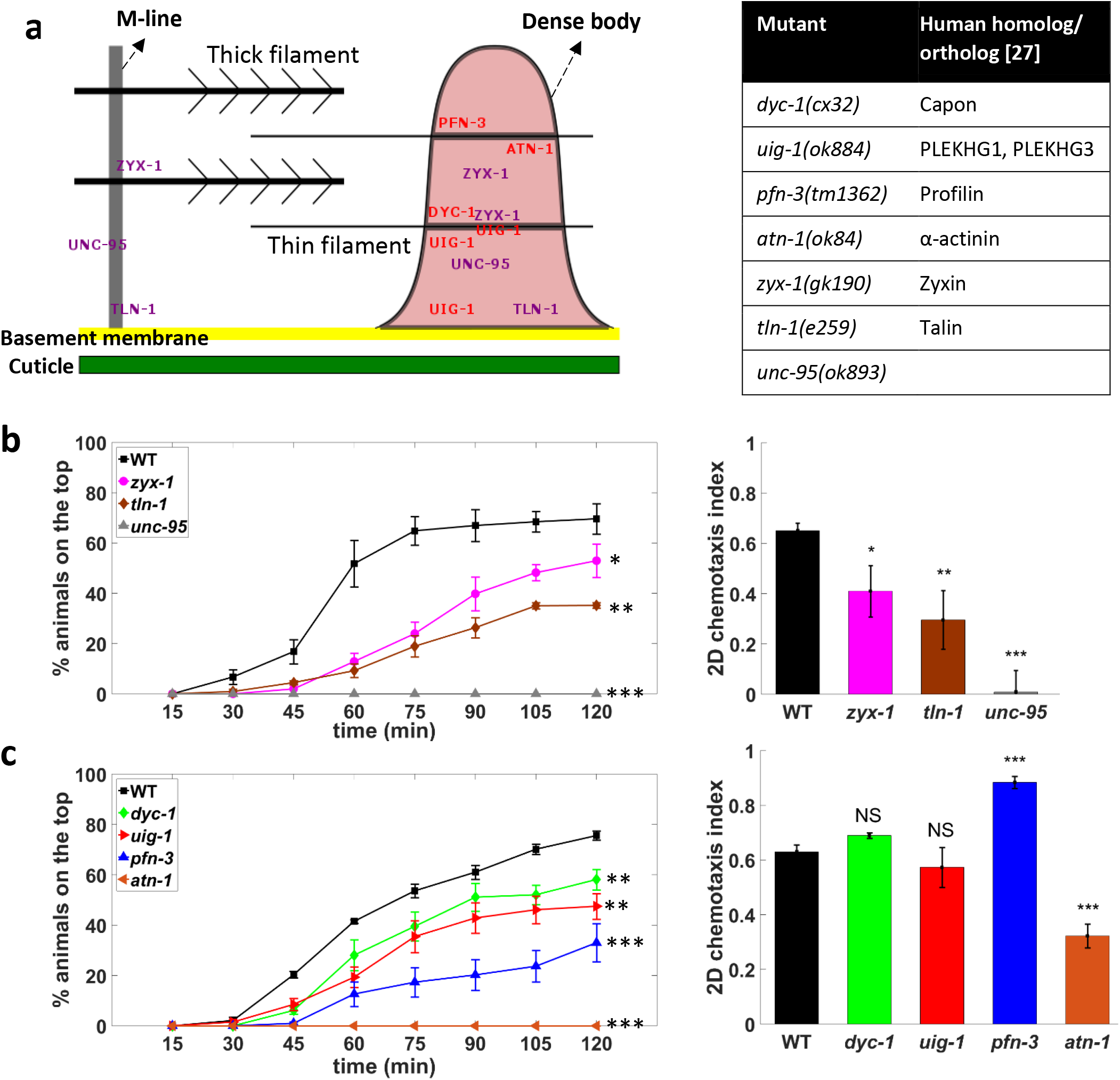
Burrowing can distinguish mutants with muscle defects. (a) Schematic showing the *C. elegans* muscle proteins that were tested, with their sites of action being on dense bodies and/or M-line. The corresponding genetic mutation and human ortholog are shown in the table. Burrowing performance and chemotactic scoring of mutants with defects in (b) both dense body and M-line, 3 replicates and (c) only dense body, 6 replicates. N = 36, 33, 34, 37, 34, 34, 35 and 34 animals for WT, *zyx-1*, *unc-95*, *tln-1*, *dyc-1*, *pfn-3*, *uig-1* and *atn-1* respectively. For burrowing, the assay conditions are 26 % w/w PF-127, H = 0.76 cm and 100 mg/mL *E. coli*. P > 0.05, *P ≤ 0.05, **P ≤ 0.01, ***P ≤ 0.001.

We tested the effect of four mutations that specifically affect the dense body (*pfn-3*, *atn-1*, *uig- 1*, *dyc-1*) and three that affect both the dense body and the M-lines (*zyx-1*, *unc-95, tln-1*). Despite these proteins localizing to the same multi-protein complexes, previously reported phenotypes were often distinct (Table S1), perhaps indicating specific functions of individual proteins or differences in the extent of the effect of the specific mutation studied. As chemotaxis is an important element of our burrowing assay, we conducted standard 2D chemotaxis assays on agar plates in parallel to ascertain if the phenotypic score of some mutants is more obvious in the burrowing assay as compared to the standard chemotaxis assay.

For the three mutants with defects in both dense body and M-line, we observed decreased burrowing performance compared with wild-type animals (Fig. 6b, left). Specifically, *unc-95* performed the worst, followed by *tln-1* and *zyx-1*. This trend in burrowing mirrored the trend we observed in 2D chemotaxis assays (Fig. 6b, right). This concordance in outcomes from 2D and 3D chemotaxis reveal that these mutants have locomotory defects, which are documented equally using either assay.

When we tested mutants with defects in dense bodies only, we found that these mutants also performed worse than wild-type with their ranking in terms of burrowing outcome decreasing as *dyc-1* > *uig-1* > *pfn-3* > *atn-1* (Fig. 6c, left). Interestingly, we found incongruence between 2D and 3D chemotaxis outcomes for some of the mutants in this cohort. *dyc-1* and *uig-1* did not show difference in 2D chemotaxis indices compared to wild-type animals, but performed poorly in burrowing (Fig. 6c, right). *pfn-3* showed a higher chemotaxis index compared with wild-type but failed to burrow effectively. Thus, while chemotaxis is an essential component of the burrowing assay, our results show that the mechanical resistance offered during burrowing can reveal differences in muscle mutants that are difficult to discern from standard 2D chemotaxis assays on agar plates.

#### ii. Modulating assay conditions to phenotype a mitochondrial mutant

Fixed assay conditions might miss the dynamic range of difference for some strains. However, our assay offers the flexibility of varying the gel height as well as concentration to increase the depth of phenotypic analysis. Here, we subjected *gei-8(gk693)* mutants to different assay conditions to reveal phenotypic differences from wild-type animals. *gei-8* encodes a homolog of the vertebrate nuclear receptor corepressor NCoR, known as a negative regulator of mitochondrial biogenesis [38]. Skeletal muscle-specific knock-out of NCoR1 in mice increased muscle mass and mitochondrial number and activity [38].RNAi-mediated knock-down of *gei-8* in *C. elegans* increased muscle mitochondrial load and oxygen consumption rate [38]. *gei- 8(gk693)* animals have a 1 kb deletion in the *gei-8* promoter region and, therefore, the potential for a modified *gei-8* expression pattern.

Our burrowing assay with 26 % w/w PF-127 and two distinct gel heights (0.7 and 0.88 cm) revealed no significant difference between *gei-8(gk693)* and wild-type animals (Fig. 7). Given that wild-type animals have the minimum burrowing ability at 30 % w/w gel concentration (Fig. 2a), we were interested in seeing if *gei-8(gk693)* would behave similarly to wild-type or manifest a different response to increased gel stiffness. Interestingly, increasing the gel concentration to 30 % w/w drops the burrowing percentage of wild type to 50% at the 2hr timepoint, while it remained around 80% for *gei-8* (Fig. 7), suggesting that *gei-8* displays superior locomotory prowess than wild-type when challenged. Although consequences of the *gei-8(gk693)* mutation on muscle strength and/or endurance remain to be clarified, and the biology of NcoR1 is likely to have complex physiology, the important aspect of the burrowing assay we highlight is that tuning the assay conditions can reveal novel and additional information about muscle function and animal behavior.

**Figure 7.**
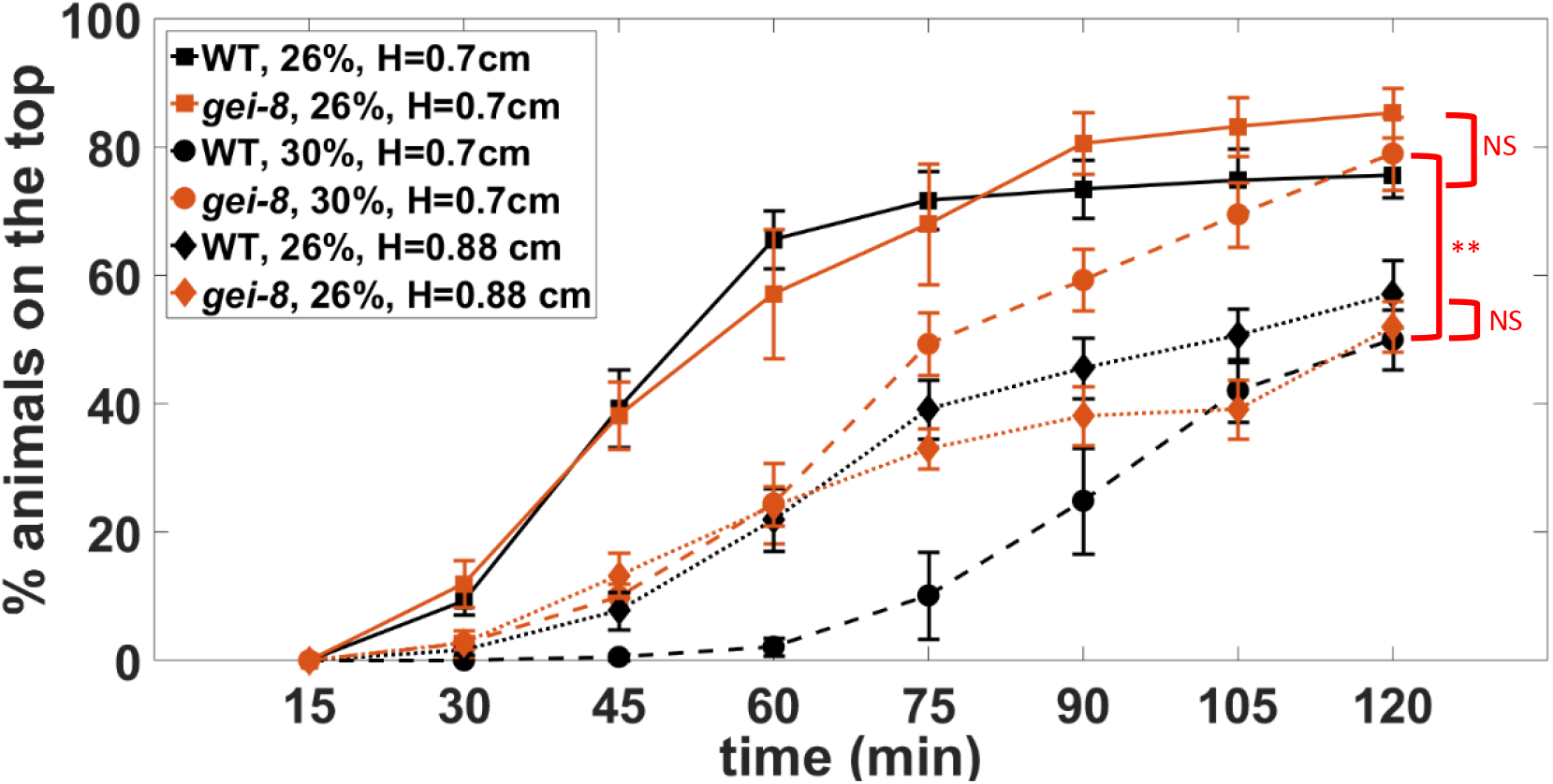
Tuning the burrowing assay conditions can reveal phenotypic differences. Burrowing performance of wild type and *gei-8(gk693)* in different assay conditions. N = 31 and 32 for wild-type and *gei-8(gk693)* respectively. Data was collected from 6 replicates. **P ≤ 0.01.

#### iii. Sorting and recovery of sub-populations for downstream molecular analysis

One of the interesting features of the burrowing assay is that not all the animals reach the attractant at the same time, i.e. some individuals burrow quickly compared with others in the population. Since animals remain at the top once they reach the food source, the Pluronic-based burrowing assay provides a simple way of sorting and recovering sub-populations of animals for downstream molecular analysis. Here, we harness this feature of the assay and show that gene expression analysis can be conducted on the fast burrowers compared with the slower ones.

We tested wild-type burrowers that reached the attractant the quickest compared with animals that were slower in burrowing and did not reach the surface within two hours (Fig. 8). We determined whether the faster burrower animals might have some differential muscle gene expression that facilitates faster movement to the surface. We selected ten body wall muscle genes known to be downregulated in expression as the animal ages [39], including troponin (*unc-27*, *tnt-2, mup-2*), tropomyosin (*lev-11*), paramyosin (*unc-15*), titin (*zig-12*), calponin (*unc- 87*), muscle myosin regulatory light chains (*mlc-1*) and myosin heavy chains (*unc-54*, *myo-3*) (see Table S2 for additional description of these genes).

**Figure 8.**
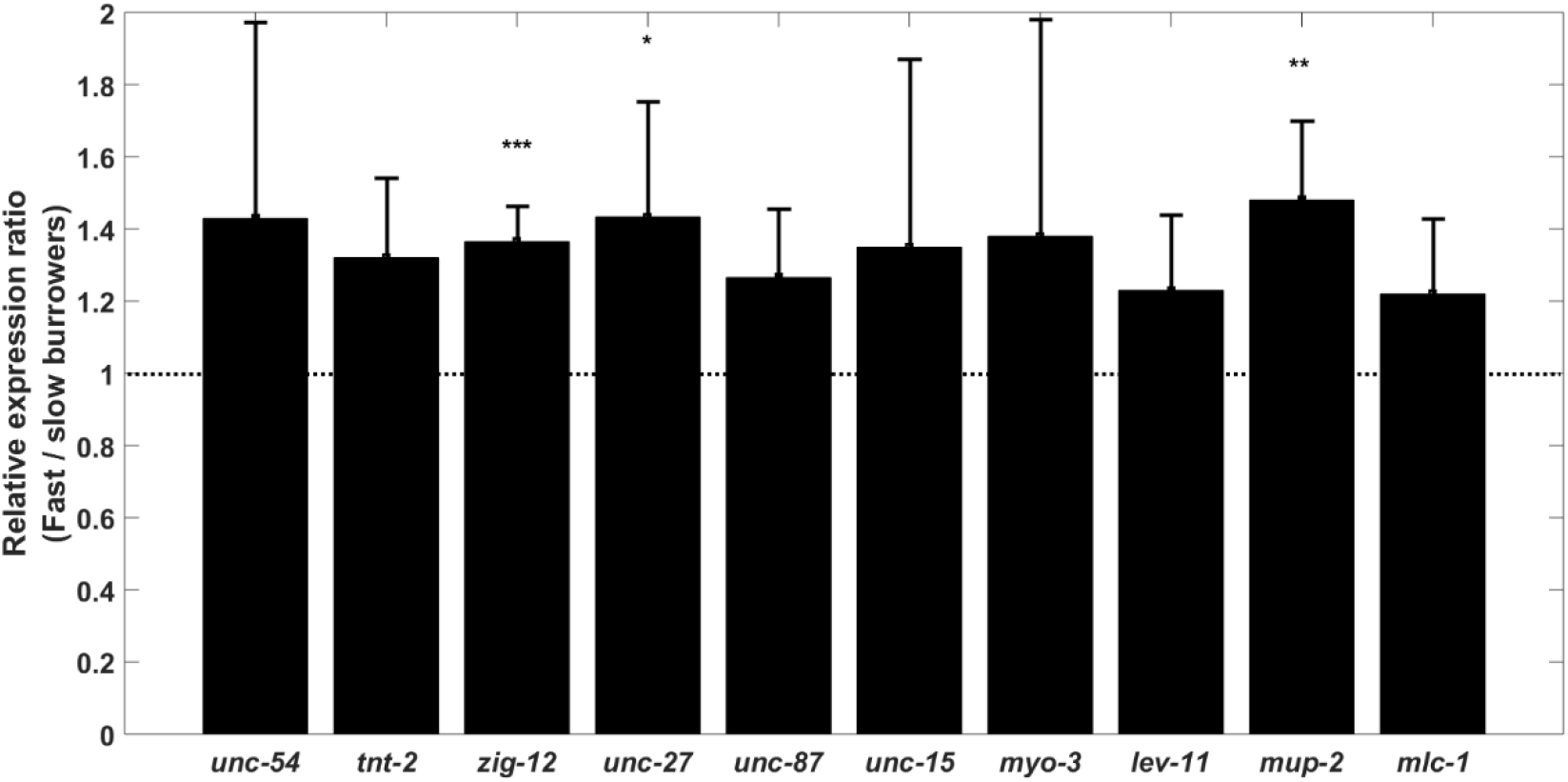
Muscle-specific gene expression in fast burrowers compared to slow burrowers. Normalized gene expression scores are shown as ratios for the successful burrowing class (among the top 10-15% that reached the top faster); to the levels for slower animals that remained in gel by the end of two-hour time period. *zig-12, unc-27, mup-2* were significantly more expressed for fast burrower animals in comparison to slower ones. Experiments were conducted with Day 1 adults. N = 30 animals with three replicates. Error bars are standard deviation. *P ≤ 0.05, **P ≤ 0.01, ***P ≤ 0.001.

We find that *zig-12, unc-27* and *mup-2* were expressed at significantly higher levels in faster burrowers (Fig.8) and the remaining genes did not show significant expression differences. However, there was a consistent upward trend for all 10 genes studied. These findings indicate the natural variation within a genetically identical population and holds potential to indicate the molecular components that might improve burrowing efficiency and locomotory prowess.

#### iv. Testing disease models of protein aggregation

*C. elegans* is widely used to model human diseases [1] including muscular dystrophies [5, 40] and neurodegenerative disorders [6]. Here, we show the utility of our burrowing assay in phenotyping *C. elegans* strains that model proteotoxicity in neurodegenerative disorders, specifically Huntington and Parkinson’s diseases. Polyglutamine expansions cause Huntington disease [41], while α-synuclein is the key protein implicated in Parkinson’s disease [42], both of which can result in toxic aggregates disrupting cellular function. Here, we focused on evaluating burrowing performance of *C. elegans* strains expressing either polyglutamine (polyQ) or α-synuclein, both fused with yellow fluorescent protein in either the body wall muscles or neurons. The polyQ strains Q35 and Q40 were expressed under the myosin promoter [43], and Q67 and Q86 were expressed in neurons [44]. For the Parkinson’s disease model, human α-synuclein protein was expressed under the myosin promoter [45].

The results from the burrowing experiments, which were conducted with day 1 adults are shown in Fig. 9. For the transgenic lines that expressed polyQ in muscle, we found that compared to wild-type, Q35-expressing animals were not deficient in their burrowing while Q40-expressing animals were significantly deficient (Fig. 9a). This difference could be due to the presence of visible protein aggregates in Q40-expressing animals and not in Q35-expressing animals (Fig. 9b). α-synuclein-expressing animals were even more impaired in their burrowing ability, as only ≈ 16 % of the animals reached the surface.

**Figure 9.**
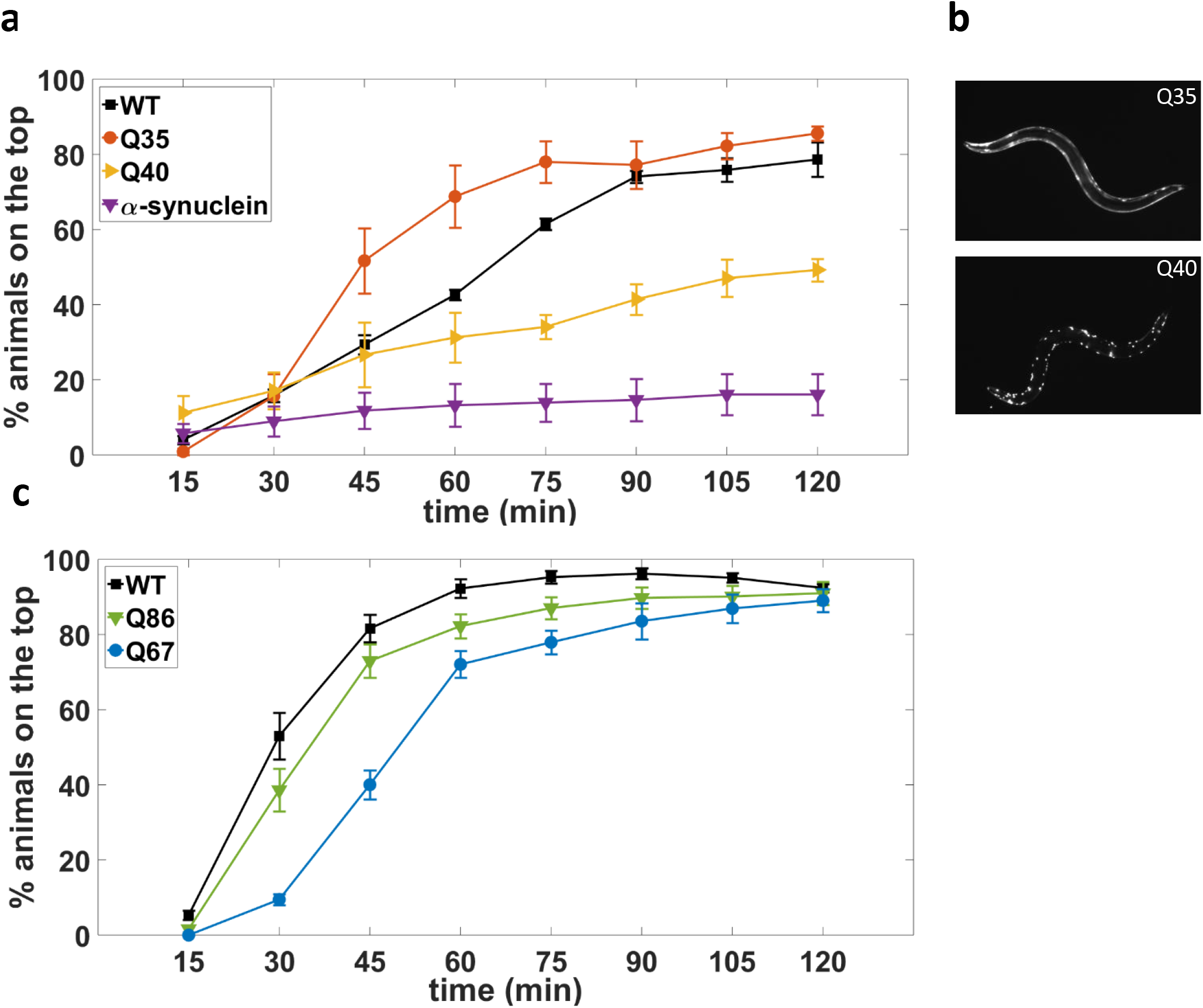
Burrowing assay phenotypes *C. elegans* models of neurodegenerative diseases. (a) Burrowing performance of day 1 adults polyQ strains with glutamine expansions in muscles, α-synuclein-expressing strain and wild-type animals. P-values are 0.1268, 0.0302 and 0.0016 for strains expressing Q35, Q40 and α-synuclein respectively. N= 34, 39, 41 and 43 animals for WT, Q35, Q40 and α-synuclein strains respectively (b) Images show visible protein aggregates in Q40-expressing strain and not in Q35-expressing strain. (c) Day 1 adults Q67-expressing and Q86-experssing animals with protein aggregation in neurons are deficient in burrowing. P-values for Q67: <0.0001, Q86: 0.0437. N= 37, 37 and 38 animals for WT, Q67 and Q86 strains respectively. Assay conditions are 26 % w/w, H = 0.7 cm and 100 mg/mL *E. coli*.

For the transgenic lines that expressed polyQ in neurons, Q67- and Q86-expressing animals were both distinctly impaired in their burrowing abilities compared with wild-type (Fig. 9c). However, the difference between wild type and Q86 was modest, and the primary differentiating factor between wild type and Q67 was the delayed rate at which Q67-expressing animals reached the surface. The burrowing impairment in transgenic *C. elegans* expressing proteotoxic aggregates in neurons therefore does not appear to be as strong as those expressed in body wall muscle, at least at the beginning of adulthood.

Our results indicate that the Pluronic-gel based burrowing assays can be used to assess declines in neuromuscular health in proteotoxic disease models of *C. elegans*, in addition to, or in place of, standard assays including thrashing and crawling [46]. For many of these strains expressing proteotoxic aggregates, motility defects have previously been reported. Q35- and Q40-expressing animals have impaired motility, with crawling defects first becoming apparent on approximately day 2 and day 0 of adulthood, respectively [43]. Thrashing defects have been reported for both Q67 and Q86 on the first day of adulthood [44]. However, day 5 animals expressing α-synuclein were not significantly worse in thrashing than their wild-type counterparts [47]. Thus, the burrowing assay has the capacity to detect neuromuscular deficits in neurodegenerative disease models and in some cases prior to major impairment, underscoring its importance for deciphering factors that influence the earliest events in degenerative processes.

## III. Discussion

### Pluronic as a novel medium for *C. elegans* burrowing

Even though burrowing of nematodes in soil-like environments has been previously studied [48–50], only recently has it been used for evaluation of neuromuscular health in *C. elegans* [16, 18, 19, 22, 51]. These recent investigations employed agar as a medium for burrowing. Here, we have shown that the biocompatible and optically transparent Pluronic gel offers an alternative for burrowing studies with several advantages including ease of manipulation of animals, flexibility in changing assay conditions to modulate burrowing performance, parallel evaluation of assay conditions or different strains in multi-well plates, and easy recovery of animals for follow-up analysis.

Our results show that the mechanical resistance offered by the Pluronic gel for burrowing is significant. Under standard assay conditions of 26 % w/w, we observed that animals burrow with a mean velocity of 1.14 mm/min and undulatory frequency of 0.07 Hz. In contrast, at 3 % w/v agar, animal velocity was found to be 2 mm/min [19] and at 9 % w/v agar the undulatory frequency was reported to be 0.2 Hz [16]. Additionally, 3D behavioral studies in 3 % w/v gelatin report a velocity of 5 mm/min [17]. Even though systematic studies comparing locomotory prowess in these different media have not been conducted, available findings indicate that the Pluronic medium is likely to be a more demanding burrowing environment for *C. elegans* than agar or gelatin.

The simplicity of our burrowing assay workflow lends itself to considerable throughput and versatility. Currently, using 12 well-plates we perform 4 strains×3 replicates in about 4 hrs that includes loading of 30 animals and the gel layer in each well, and scoring every 15 mins for a total duration of 2 hrs. Other assay formats could also be designed, where for example, the fraction of animals on the gel surface is recorded only at the final time point. Such end-point assays can be performed with significantly higher throughput and in a miniaturized format involving 96 well-plates.

### Burrowing is a sensitive phenotyping tool to evaluate neuromuscular function in *C. elegans*

Very few studies have exploited burrowing ability as a way to evaluate neuromuscular health in *C. elegans*. Since the original method was conceived [18], it has been applied to score muscular dystrophy mutants [16], and to probe calcium dysregulation and longevity outcomes due to physical exertion in the burrowing environment in these mutants [19]. Additionally, a burrowing assay has been used to assess mutants with defects in mitochondrial fission and fusion proteins [22]. Here, we have developed an improved burrowing method based on a thermoreversible Pluronic gel and expanded its application to diverse areas with novel findings.

Our results show that mutants can be uniquely distinguished based on burrowing capacity, which is sometimes difficult to discern from 2D crawling locomotion and thrashing assays. As shown in Table S1, the muscle mutants studied in Sec. II.D-i could not be differentiated based on just one phenotypic assay suggesting that subtle genetic defects might be difficult to detect by existing locomotory assays. The fact that burrowing distinguishes all the muscle mutants studied indicates its capacity to detect subtle defects in neuromuscular function.

In general, successful burrowing of *C. elegans* requires that both neuromuscular function and chemotaxis responses must be intact. How do we dissect the contributions of neuromuscular strength and chemotactic prowess from burrowing experiments? Using 2D chemotaxis in conjunction with burrowing can help address these contributions to a certain extent as evident from the muscle mutants in Fig. 6c, which do not show obvious 2D chemotaxis deficits but exhibit significant burrowing deficiency. In cases, where mutants show impairment in both 2D chemotaxis and burrowing, additional assays can be utilized. For example, microfluidic systems can be used to score for muscle strength in *C. elegans* [9] based on deflectable micropillars or chemotactic ability can be recorded from head-swaying of partially-immobilized animals, minimizing the contribution of crawling locomotion [52].

Our burrowing platform offers the potential for several new lines of investigation. For example, the muscle mutants *atn-1* and *unc-95* hardly burrow toward the attractant, indicating that mutagenizing these strains and conducting our assays might reveal compensatory mechanisms that may promote burrowing. Likewise, transcriptomic analysis on the fast burrowers is likely to uncover new genes that improve burrowing ability and therefore neuromuscular health in *C. elegans*. Our results on Huntington and Parkinson’s disease models (Fig. 9) certainly opens the door for screening of compounds that can lead to prioritization of therapeutic candidates for testing in mammalian systems.

## Conclusions

We have shown that configuring the *C. elegans* burrowing assay in Pluronic gel medium offers the convenience of a simple workflow with high throughput. We systematically studied the influence of assay conditions on burrowing ability and modulated the physical challenge experienced by animals based on the needs of a particular application. The demonstrative applications we have chosen highlights the richness of the burrowing assay in diverse areas where the neuromuscular system is implicated. We anticipate that the Pluronic burrowing assay can be easily adopted in other laboratories, enabling comprehensive investigations of the molecular, cellular and tissue-level mechanisms required for the maintenance of neuromuscular health in *C. elegans* that might be translatable to human neuromuscular diseases.

## Materials and methods

### Worm culture

*Caenorhabditis elegans* wild type N2 was cultured at 20 °C on standard nematode growth medium (NGM) on 60 cm petri plates and never allowed to starve. The NGM plates were allowed to dry 24 hours prior to seeding with 450-550 μL of *Escherichia coli* OP50 bacteria overnight. Age synchronization was done by transferring 20-25 gravid animals to seeded plates and letting them lay eggs for 3-4 hours. After the desired number of eggs were laid, the gravid adults were removed, and the eggs were allowed to hatch and develop in the 20 °C incubator. The animals used for all experiments were day 1 adults. The day that the age synchronized animals started to lay eggs is counted as day 0 of adulthood.

The following mutants were obtained from *Caenorhabditis* Genetics Center: *dyc-1(cx32), pfn-3(tm1362), uig-1(ok884), atn-1(ok84), zyx-1(gk190), unc-95(ok893), tln-1(e259).* For calcium imaging the following strain was used - HBR4: goeIs3 HBR4: goeIs3[*Pmyo-3*::GCaMP3.35::*unc-54*-3’utr, *unc-119*] which expresses the calcium indicator GCaMP3 in body wall muscles. The strains were imaged using a Nikon Ti-E microscope at 15× with an exposure of 150 ms, emission wavelength of 535 nm, and excitation wavelength of 480 nm.

Strains used as protein aggregation disease models have the following genotypes: pkIs2386[*Punc-54*::alphasynuclein::YFP + *unc-119(+)*], rmIs132[*Punc-54* Q35::YFP], rmIs133[*Punc-54* Q40::YFP], N2; rmEx135(F25B3.3p::Q86::YFP), and N2; rmEx164(F25B3.3p::Q67::YFP).

### Pluronic-based burrowing assays

To develop and optimize the burrowing assay, various Pluronic concentrations and gel heights were tested. Pluronic F127 (Sigma-Aldrich) was dissolved in deionized water to reach the desired concentration, reported as solute weight per solution weight (% w/w). The suspension was kept at 4 °C until all the Pluronic pellets dissolved. The solutions were stored at 4 °C prior to the experiment to prevent gelation. The Pluronic concentration in all assays was the optimized concentration of 26 % w/w, except for when other concentrations were tested.

The Pluronic solutions were kept at 14 °C in a Waverly digital thermal bath for at least 30 minutes before conducting the experiment. 20-30 μL of solution was added to the bottom of a Corning™ Falcon™ Polystyrene 12-well plate (Fig. 1a). A minimum number of 30 animals (day 1 adults) were hand-picked from NGM plates and released into the drop. After around 10 minutes, a layer of Pluronic was cast on top of the initial droplet, to the desired thickness ranging from 0.44 cm to 1.1 cm depending on the experiment. The top layer needed around 5 minutes to gel at a room temperature of 20 °C. Gelation was confirmed by no fluid movement and inserting the tip of the worm pick somewhere close to the edge of the well. If no healing of the indentation was observed after piercing, then the gel had been formed, and 20 μL of the chemoattractant was added directly to the top (t=0 min). The animals that had burrowed to the surface were scored by monitoring the top layer under the microscope and counting the ones that had reached the top every 15 minutes for a total duration of 2 hours. The replicates per treatment, each in a different well, were conducted concurrently. The percentage of animals on the top surface was defined as the number of animals on the top surface divided by the total sample size in that well.

All the burrowing assays were conducted by hand picking animals, except Fig. 9a on neurodegenerative disease models, as those animals were rinsed off the plates with DI water and collected in Falcon tubes. The rinsing process is the same as preparing animals for chemotaxis assays (See Methods on Chemotaxis assays). After rinsing off the animals from bacteria, 10 µL of worm solution was placed on the bottom of well plate. Then, 500 µL of PF-127 was added to make a base gel layer. Finally, the Pluronic layer was cast on top to the desired thickness of 0.7 cm, followed by 20 µL of *E. coli* attractant.

### Behavioral analysis during burrowing

The animals were assessed for burrowing behavioral analysis and characterization by being sandwiched in 1 mL Pluronic gel between a 75 mm × 25 mm Fisherbrand™ plain microscope glass slide and a coverslip with a gel thickness of approximately 1 mm. This method was utilized to reduce the chance of a vertically oriented planar burrowing rather than parallel to the microscope focal plane for the sake of better imaging. After sandwiching, the animals were allowed 30 minutes to acclimate before the imaging started. For velocity and undulatory frequency, 2-minute video segments in which the animals were exclusively moving forward were selected. Eight 15-second unique time segments were the replicates used to find the velocity. The worm body centroid was tracked and the distance the centroid traveled every 15 seconds was measured using ImageJ software [53]. The undulatory frequency (Hz) was defined as the number of full-wavelength sinusoidal movements the worm could make within one second. For this purpose, the required time for a full sinusoidal movement was measured for eight distinct sinusoidal movements. Then, the inverse was taken to result in the frequency (Hz).

To characterize the burrowing behavior, 5-minute uninterrupted videos with the frame rate of 3 fps were captured using a Nikon Ti-E microscope at 4×. Afterwards, the videos were analyzed by assessing the time the animals spent moving forward, backward, or pausing.

To investigate animal distribution across gel heights, the total gel height was divided into three equal layers, and the number of animals in each segment at each time point was counted by scanning the z-axis through the gel height from bottom to the top surface using a Nikon Ti-E microscope with a 4× objective lens.

### 2D Chemotaxis assays

To test the effectiveness of animal chemosensation, standard 2D chemotaxis assays were conducted. *E. coli* was concentrated in liquid NGM (as above without agar). Isoamyl alcohol and diacetyl were diluted in ethanol to 1%. For agar chemotaxis experiments, the chemoattractant solutions were freshly mixed with an equal volume of 0.5 M sodium azide as an anesthetic, facilitating scoring.

Volatile compound chemotaxis assays on agar plates were conducted according to Margie *et al.* [54] with a few modifications. Briefly, chemotaxis agar (2% agar (Fisher Scientific), 5 mM potassium phosphate (Fisher Scientific), 1 mM calcium chloride (Avantor), 1 mM magnesium sulfate (Sigma-Aldrich)) was prepared a day prior to the experiment and poured onto 6 cm petri plates. The animals were collected in Falcon tubes by washing them from culture plates with deionized water and allowing them to pellet by gravity. After 5 minutes, the supernatant was carefully removed, water was added, and the tube was inverted a few times to wash the worms. This rinsing process was repeated 4 times, so the supernatant on top looked clear and bacteria-free. Finally, the animal pellet was resuspended in water to obtain around 60 animals per 5 μL. 5 μL of worm solution was pipetted onto the center of the chemotaxis plate. Immediately, 2 μL of test solution (chemoattractant) and 2 μL of control solution (diluent) were added to their designated quadrants. After an hour, chemotaxis indices were calculated as the difference in the number of worms in the test and control quadrants (excluding the animals that did not migrate farther than 1 cm), normalized with the total number of worms.

Chemotaxis assays with *E. coli* in liquid NGM required time for gradient formation [55] and was conducted according to Wen *et al.* with minor modifications [56]. The chemotaxis agar was made as described in the volatile compounds assay section above. 10 mL of chemotaxis agar was poured into 10 cm petri plates a day prior to the experiment. The plates were marked underside by dividing them into two semicircles. On one side, the test solution (5 μL of *E. coli* in liquid NGM) was spotted 3 cm from the midline. The control solution (5 μL of DI water) was placed on the exact opposite side. After the solutions were placed, the plates were left for 3.5 hours before testing the animals to allow the chemical gradient to establish. The animals were washed from their culture plates with chemotaxis buffer (5 mM potassium phosphate, 1 mM calcium chloride, 1 mM magnesium sulfate) until they were free of bacteria. The total time that the worms spent swimming in liquid was kept under 30 minutes to minimize fatigue [57]. Then, 5 μL of the solution containing animals was added onto the center. Whenever the liquid absorbed, the animals were gently dispersed using a worm pick, so they were not trapped within the liquid droplet. The assay was completed in 1.5 hours, and the chemotaxis index was calculated as mentioned before.

### RNA extraction and quantitative PCR

RNA extraction and quantitative PCR were carried out as previously described [57]. Briefly, we collected N2 *C. elegans* after burrowing (~30 animals per sample) into TRIzol Reagent (Ambion) and immediately froze animals in liquid nitrogen. The quick burrowers were among the top 10-15% in burrowing performance, while the slow burrowers were recovered from the Pluronic at the conclusion of the two-hour time period. After freeze-thaw cycles with liquid nitrogen/37 °C heat block, we extracted total RNA following the manufacturer’s instructions (Ambion) and synthesized cDNA using the SuperScript III First-Strand Synthesis System (Invitrogen).

We performed quantitative PCR using diluted cDNA, PerfeCTa SYBR Green FastMix (Quantabio) and 0.5 µM of gene-specific primers (Table. S3) in a 7500 Fast Real-Time PCR System (Applied Biosystems), calculating relative expression using the ΔΔCt method [58] with *cdc-42* and Y45F10D.4 as reference genes [59].

### Statistical analysis

To compare mutants and disease models burrowing performance, two-way ANOVA was used in GraphPad Prism software. The statistical analysis on chemotaxis indices was done using two-sample student’s *t*-test in MATLAB. For qPCR, paired-sample *t*-test was done using MATLAB. Burrowing assays error bars represent standard error of the mean. All the assays were done in three replicates unless otherwise noted. N represents average of the number of animals used in the replicates.

## Supporting information

Video S1. 2-hr time-lapse of burrowing assay

Video S2. Pmyo-3 GCaMP3.35 burrowing in 26 % w/w PF-127

## Acknowledgements

Some strains were provided by the CGC, which is funded by NIH Office of Research Infrastructure Programs (P40 OD010440). We would like to thank Anam Mahmood for assistance with experiments and Guy M. Benian for useful discussions. This work is partially supported by funding from the National Institutes of Health (RO1 AG051995-04 to MD & SV), Cancer Prevention and Research Institute of Texas (RP160806 to SV), National Aeronautics and Space Administration (NNX15AL16G to SV & JB) and the Biotechnology and Biological Sciences Research Council (BB/N015894/1 to NJS). JEH acknowledges funding support from Fulbright scholar program. RL has been funded by postdoctoral fellowships from Life Sciences Research Foundation (sponsored by Simons Foundation) (award # Laranjeiro-2015) and American Heart Association (award # 18POST33960502). AA was supported by the Max Planck Society.

## Author contributions

LL, CMRL and SAV conceived the Pluronic gel-based burrowing assay. LL, JEH and RL performed the experiments. All authors analyzed and interpreted the data. LL, JEH and SAV wrote the paper. All authors read and commented on the manuscript. SAV supervised the study.

