## Supplementary information for "Pluronic gel-based burrowing assay for rapid assessment of neuromuscular health in *C. elegans*"

Supplementary Video S1. 2-hr time-lapse of burrowing assay.

Supplementary Video S2. Pmyo-3 GCaMP3.35 burrowing in 26 % w/w PF-127.

Supplementary Figure 1. Testing the effect of gravity on burrowing in 26 % w/w Pluronic

Supplementary Figure 2. Pluronic elastic modulus as a function of shear stress and Pluronic concentration

Supplementary Figure 3. Characterization of burrowing behavior in 26 % w/w PF-127

Supplementary Table 1. Dense body and M-line mutants used to study the muscle defects

Supplementary Table 2. List of genes tested for qPCR

Supplementary Table 3. List of primers used for qPCR

**Supplementary Video S1. 2-hr time-lapse of burrowing assay.** Wild-type animals are burrowing toward the attractant (*E. coli*) on the top surface in 26 % w/w PF-127 within the assay duration of 2 hours.

**Supplementary Video S2. Pmyo-3 GCaMP3.35 burrowing in 26 % w/w PF-127.** As muscle actuates during burrowing, they appear brighter due to an elevation in calcium ions levels. In the beginning, the animal has acquired a 3D posture, so not all the body appears in the same focal plane. As it moves forward, the tail is emerging in the same focal plane of the microscope, making all the animals body get in focus.

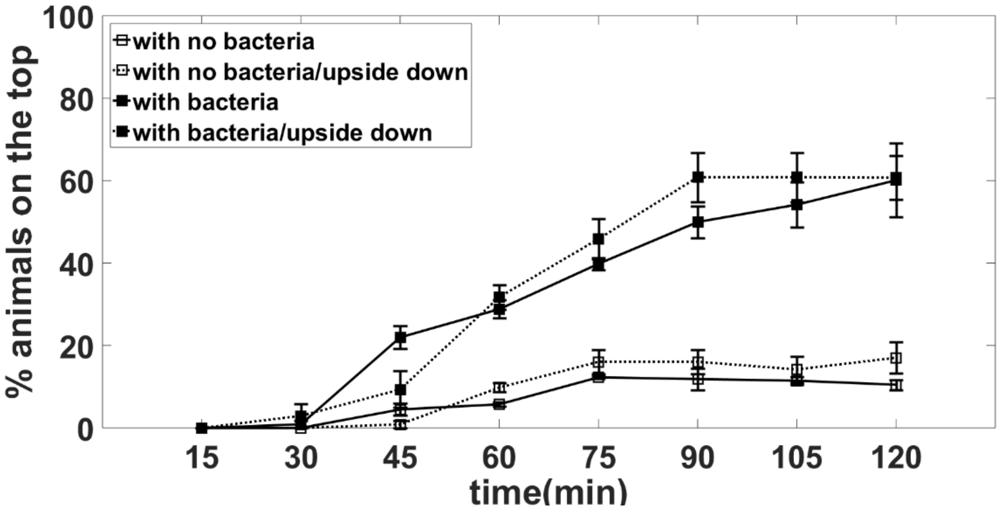


**Figure S1. Testing the effect of gravity on burrowing in 26 % w/w Pluronic**. Gravity did not show any significant effect on the burrowing rate of wild-type animals. Gel thickness, H = 0.9 cm (N= 37 on average. 3 replicates per condition).

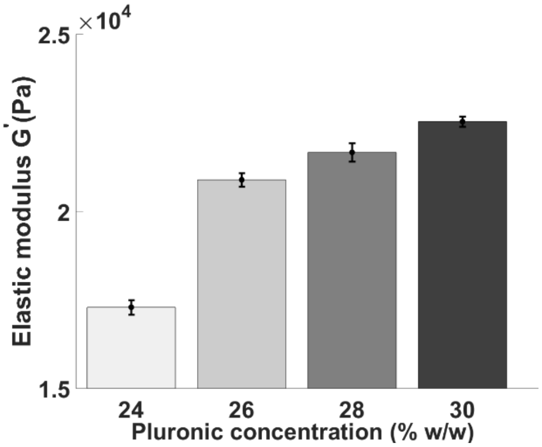


| Pluronic concentration (% w/w) | Yield stress (Pa) |
|---|---|
| 24 | 251.2 |
| 26 | 316.2 |
| 28 | 398.1 |
| 30 | 501.2 |

**Figure S2. Gel elastic modulus and yield stress as a function of Pluronic concentration**.
The elastic modulus for 24 % w/w 26 % w/w, 28 % w/w and 30 % w/w are 17.3 kPa, 20.9 kPa, 21.7 kPa and 22.5 kPa, respectively. The yield stress for 24 % w/w to 30 % w/w varies from 251.2 Pa to 501.2 Pa.
Rheology measurements were performed using an AR2000 rheometer with parallel plate geometry (40 mm diameter and 500 $\mu$m gap). The base plate was initially set at 10 ˚C and equilibrated at 20 ˚C for doing all the measurements. Samples were kept at 4 ˚C prior to the experiment. Oscillatory shear stress sweep from 1 to 1000 Pa at the frequency of 1 Hz was used to find the linear viscoelastic region. The yield stress was determined as the critical oscillatory stress when the elastic modulus deviated more than 10% from the linear viscoelastic region.

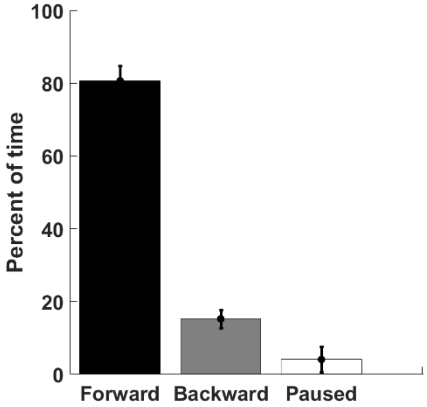


**Figure S3. Characterization of burrowing behavior in 26 % w/w PF-127.** Percent of time duration wild-type animals spent in each phase of locomotion. Animals spent most of their time moving forward.

**Table S1. Dense body and M-line mutants used to study the muscle defects**

| Mutant | Human homolog/ ortholog [1] | Phenotype |
| --- | --- | --- |
| *dyc-1(cx32)* | Capon | Hyperactivity, overbent [2] |
| *uig-1(ok884)* | PLEKHG1, PLEKHG3 | Normal swimming frequency, Higher crawling frequency, higher minimum radius of curvature [3], normal bending [4] |
| *pfn-3(tm1362)* | Profilin | Slightly decreased thrashing [5], bending-defective [4] |
| *atn-1(ok84)* | α-actinin | Bending-defective [4], abnormal dense bodies, normal thrashing [6], higher minimum radius of curvature, normal crawling [3] |
| *zyx-1(gk190)* | Zyxin | Bending-defective [4] |
| *tln-1(e259)* | Talin | Unc phenotype [7], exaggerated body bends (www.wormbase.org) |
| *unc-95(ok893)* | | Movement defect [8], bending-defective [4], disrupted dense body and M-line [9] |

**Table S2. List of genes tested for qPCR**

| Gene | Human ortholog or homolog | Description (From WormBase WS264) |
|---|---|---|
| *unc-54* | Muscle myosin heavy chain (MHC B) | Required for locomotion and egg-laying a thick filament component that is expressed in multiple muscle cell classes |
| *tnt-2* | Troponin T | Expressed in the anal depressor muscle, reproductive system, and the body wall musculature. |
| *zig-12* | Titin | Localized to the endoplasmic reticulum, the sarcomere and the striated muscle dense body |
| *unc-27* | Troponin I | Required for coordinated motility, normal muscle morphology and proper sarcomeric organization |
| *unc-87* | Calponin like | maintain the structure of myofilaments in body wall muscle cells |
| *unc-15* | Paramyosin | Physically interacts with MHC A, one isoform of myosin heavy chain (MHC) in striated muscle |
| *myo-3* | Myosin heavy chain | Essential for thick filament formation, and for viability, movement, and embryonic elongation; expressed in body muscle |
| *lev-11* | Tropomyosin 1 | An actin-binding contractile structural protein, is required for embryonic development, normal body morphology, and locomotion |
| *mup-2* | Troponin T | Affects embryonic body wall muscle cell contraction, sarcomere organization, cell positioning, regulated muscle contraction in larval and adult body wall muscle |
| *mlc-1* | Muscle regulatory myosin light chain | Functions in the pharyngeal and body-wall muscle development, affects locomotion and growth; expressed in the body-wall muscles, pharyngeal muscles, and vulval muscles |

**Table S3. List of primers used for qPCR** (*cdc-42* and Y45F10D.4 are the control references)

| Gene | Forward primer (5'-3') | Reverse primer (5'-3') | Amplification efficiency |
|---|---|---|---|
| **cdc-42** | CTGCTGGACAGGAAGATTACG | CTCGGACATTCTCGAATGAAG | 1.03 |
| **lev-11** | CCGCTGAAGAGAAAGTCCGT | TCGTCTCCGGTCTGAGTCAT | 0.93 |
| **mlc-1** | TGGAGCCTTTGCCATGTTC | CTTGACCTCATCCTCGTCCAAT | 0.99 |
| **mup-2** | AACGCAAGGCTAAGGCTGAT | AGCTCCGGCTTCAACTCTTC | 0.92 |
| **myo-3** | AGGGAGACTTGAAGGTTGCG | AGCGAGCTTAGCATTGGTGT | 0.93 |
| **tnt-2** | ATGGGGACGCAAAGAGAACG | AATTGGGTTGACTGGTGGCT | 1.00 |
| **unc-15** | CGAGGAAGCCAATGGACGTA | AAATCAGCTTGAGCGGTGGA | 0.95 |
| **unc-27** | CGTGGAAAGTTCGTCAAGCC | TCTTGAGGTTGGCACGGAAG | 0.98 |
| **unc-54** | ACTACCAACACGAAGCCGAG | GGCGTTAGCCTTGGAGAGTT | 0.94 |
| **unc-87** | ATGACTGGATTCGGACAGCC | AGCTTGAGAAGCAAAACGGT | 0.98 |
| **Y45F10D.4** | GTCGCTTCAAATCAGTTCAGC | GTTCTTGTCAAGTGATCCGACA | 0.94 |
| **zig-12** | GATCAGAGAACGGGTCGGTG | CTCCTCAAGCTCGTCTGGTC | 1.02 |
